# Characterization of two lytic bacteriophages isolated from urban surface water in Romania targeting multidrug-resistant *Escherichia coli*

**DOI:** 10.64898/2026.05.08.723789

**Authors:** Rareş-Ionuţ Dragomir, Tudor-Emanuel Fertig, Coralia Bleotu, Mariana Carmen Chifiriuc, Ilda Czobor Barbu

**Affiliations:** The Research Institute of the University of Bucharest, ICUB, Şoseaua Panduri 90, District 5, Bucharest, 050663, Romania; Department of Botany and Microbiology, Faculty of Biology, University of Bucharest, Splaiul Independenţei 91-95, Bucharest, 050095, Romania; Biological Sciences Division, Romanian Academy, Calea Victoriei 125, Sector 1, Bucharest, 010071, Romania; Ultrastructural Pathology and Bioimaging Lab., “Victor Babeş” National Institute of Pathology, Bucharest, Romania; Cellular and Molecular Pathology Dept., “Ştefan S. Nicolau” Institute of Virology, Bucharest, Romania

## Abstract

**Background:** The global rise of multidrug-resistant (MDR) bacteria represents a critical public health threat, and Romania ranks amongst the most affected countries in Europe. As conventional therapy increasingly fails, bacteriophage therapy has re-emerged as a promising alternative to antibiotics. Urban rivers, contaminated with resistant bacterial strains, represent an underexplored and accessible reservoir for the isolation of lytic phages with therapeutic potential.

**Methods:** Two bacteriophages, 17M_Ec17_D and 22C_Ec22_D, were isolated from the Dâmboviţa River, Bucharest, Romania, using MDR *E. coli* as host bacteria. Phage characterization included plaque morphology, transmission electron microscopy, and host range assessment by spot assay against 30 MDR *E. coli* isolates. Whole genome sequencing was performed on Illumina MiSeq and Oxford Nanopore Technologies MinION platforms, followed by bioinformatic analysis including taxonomic classification, lifestyle prediction, and functional annotation.

**Results:** Both phages formed clear plaques and were classified as *Kayfunavirus* (17M_Ec17_D, *Podoviridae*-like) and *Kagunavirus* (22C_Ec22_D, *Siphoviridae*-like) with nucleotide similarities of 89.2% and 71.4% to their closest relatives, respectively, suggesting both are candidates for novel species. Host range analysis revealed lytic activity against 13% and 10% of tested MDR isolates, with complementary infection profiles. Genomic analysis confirmed a strictly lytic lifestyle for both phages, supported by the presence of holin and spanin genes and the absence of lysogenic modules, antibiotic resistance genes, and virulence factors.

**Conclusions:** To the best of our knowledge, this is the first study conducted in Romania to isolate and genomically characterize lytic bacteriophages targeting MDR *E. coli*. The characterized phages represent safe therapeutic candidates whose complementary host ranges suggest potential application as part of phage cocktail to broaden antimicrobial coverage against MDR infections.

## 1. Introduction

Antibiotic resistant bacteria (ARB) represent a major 21st-century threat, ranking among the top 10 global public health challenges. Beyond human health, ARB also pose significant risks to the food industry and environmental protection, with substantial associated economic losses. According to the most recent data, in 2021, approximately 4.71 million deaths were associated with ARB, of which more than 1.14 million were directly attributable (Naghavi et al., 2024). Romania ranks among the European countries with the highest levels of ARB, according to the European Centre for Disease Prevention and Control (ECDC, 2021). Forecast for 2050 predicts that over 39 million deaths globally could be directly linked to the ARB (Naddaf, 2024).

The rapid emergence of multidrug-resistant (MDR) bacteria represents one of the most critical challenges to modern medicine. Among these pathogens, *Escherichia coli* is particularly concerning due to its prevalence in both clinical infections and environmental reservoirs. As standard antibiotics increasingly fail to control these infections, there is an urgent need for alternative therapeutic strategies.

Bacteriophages (phages) are the most abundant biological entities of the world. They are taxonomically classified into multiple orders and families based on morphology and infection mechanisms. Of these, the *Caudovirales* order is considered the most effective against ARB. Members of this group possess an icosahedral capsid and a contractile tail structure, along with a double-stranded DNA genome. Significant advantages of phages are their high target specificity and their ability to disrupt bacterial biofilms, a common resistance mechanism, by secretion of lytic enzymes such as lysins, which degrade the extracellular matrix of biofilms (Dicks & Vermeulen, 2024; Egido et al., 2022).

Urban wastewater treatment plants (WWTPs) are known to be rich reservoirs for both resistant bacteria and the phages that prey upon them, making them ideal sites for bioprospecting new therapeutic agents (Shivaram et al., 2023).

This study aims to isolate and characterize bacteriophages from the Dâmboviţa River, Bucharest, Romania. By combining morphological observation with advanced Nanopore and Illumina sequencing, we seek to evaluate the genomic safety and lytic efficacy of these isolates against a panel of MDR *E. coli* strains.

## 2. Materials and Methods

### 2.1 Phage isolation and purification

Two bacteriophages were isolated from grab water samples collected from the Dâmboviţa River, Bucharest, Romania (44.43366° N, 26.081418° E) and filtered through a 0.45 µm PES filter. Phages were isolated using an enrichment method in which 25 MDR bacterial isolates of *Escherichia coli* and *Klebsiella pneumoniae* were incubated with filtered water and Luria-Bertani (LB) medium for 24 hours at 37°C. After the incubation period, samples were centrifuged at 12,000 × g for 5 minutes and then filtered through a 0.22 µm PES filter. Phage suspensions were serially diluted using Salt-Magnesium (SM) buffer, and plaque assays were performed to determine plaque morphology and suspension purity for each isolated phage. Each phage suspension was purified by picking single plaques at least three times. A high-titer phage lysate was obtained using the plate lysis method, where plates were flooded with 10 mL of SM buffer and incubated at 4°C for 1 hour. After the incubation period, each suspension was centrifuged at 12,000 × g for 10 minutes at 4°C and then filtered through a 0.22 µm PES filter. The plaque-forming units per milliliter (PFU/mL) were determined by plaque assay for each isolated phage. Stocks were prepared by combining phage suspension with 15% glycerol and stored at -80°C.

### 2.2 Host range

Lytic activity of isolated phages was assessed using spot assay against 30 MDR *E. coli* isolates (12 clinical and 18 environmental isolates). A bacterial lawn was prepared on LB agar plates, and three 10 µL spots of each freshly prepared phage suspension were applied and incubated for 24 h at 37°C.

### 2.3 DNA extraction and Whole Genome Sequencing (WGS)

Fresh high-titer phage suspensions were precipitated with PEG-8000 and incubated at 4°C for 24 h. Phage DNA was extracted using the phenol:chloroform:isoamyl alcohol (25:24:1) method and subsequently quantified using a NanoDrop and Qubit 3.0 fluorometer.

WGS was carried out on two platforms, Illumina MiSeq (v3, 600 cycles) using the Nextera XT Library Prep Kit (Illumina, San Diego, CA, USA) and Oxford Nanopore Technologies MinION Mk1b using the Rapid Barcoding Kit SQK-RBK114.96 (Oxford Nanopore Technologies, Oxford, England).

### 2.4 Bioinformatic Analysis

*De novo* assembly of raw reads was performed using Flye v2.9.6 (Kolmogorov et al., 2020), and the resulting FASTA files were subjected to further analysis. dsDNA phages were classified using the taxMyPhage v0.3.7 (Millard et al., 2025) tool in accordance with the ICTV database. The BACPHLIP (Hockenberry & Wilke, 2021) tool was employed to determine phage lifestyle. Whole-genome annotation was performed with Pharokka v1.9.1 (Bouras et al., 2023) using the PHROG (Terzian et al., 2021) database. Virfam online software was used for phage classification based on head-neck-tail modules (Lopes et al., 2014).

## 3. Results

### 3.1. Phage morphology and host range

The two isolated and purified phages, 17M_Ec17_D and 22C_Ec22_D, each formed clear plaques with uniform morphology. A semi-transparent halo was additionally observed for both phages (Fig. 2A, B and Fig. 3A, B) after 24 h at 37°C on overlayer agar. The titer of 17M_Ec17_D was 7.6 × 10^13^ PFU/mL and that of 22C_Ec22_D was 3.13 × 10^10^ PFU/mL. Host range analysis showed that 17M_Ec17_D exhibited lytic activity against 13% of the tested MDR isolates, and 22C_Ec22_D against 10% (Fig. 1).

**Fig. 1.**
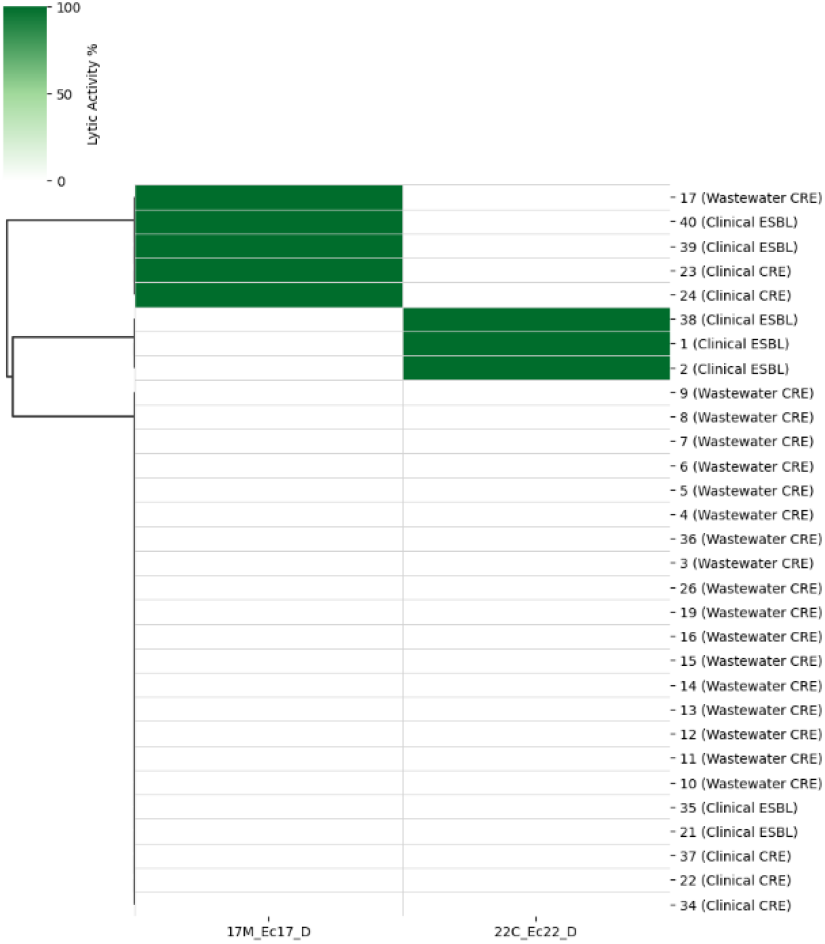
Host range of phages 17M_Ec17_D and 22C_Ec22_D. Lytic activity assessed by spot assay against 30 MDR *E. coli* isolates (12 clinical and 18 environmental). Green cells indicate lytic activity.

**Fig. 2.**
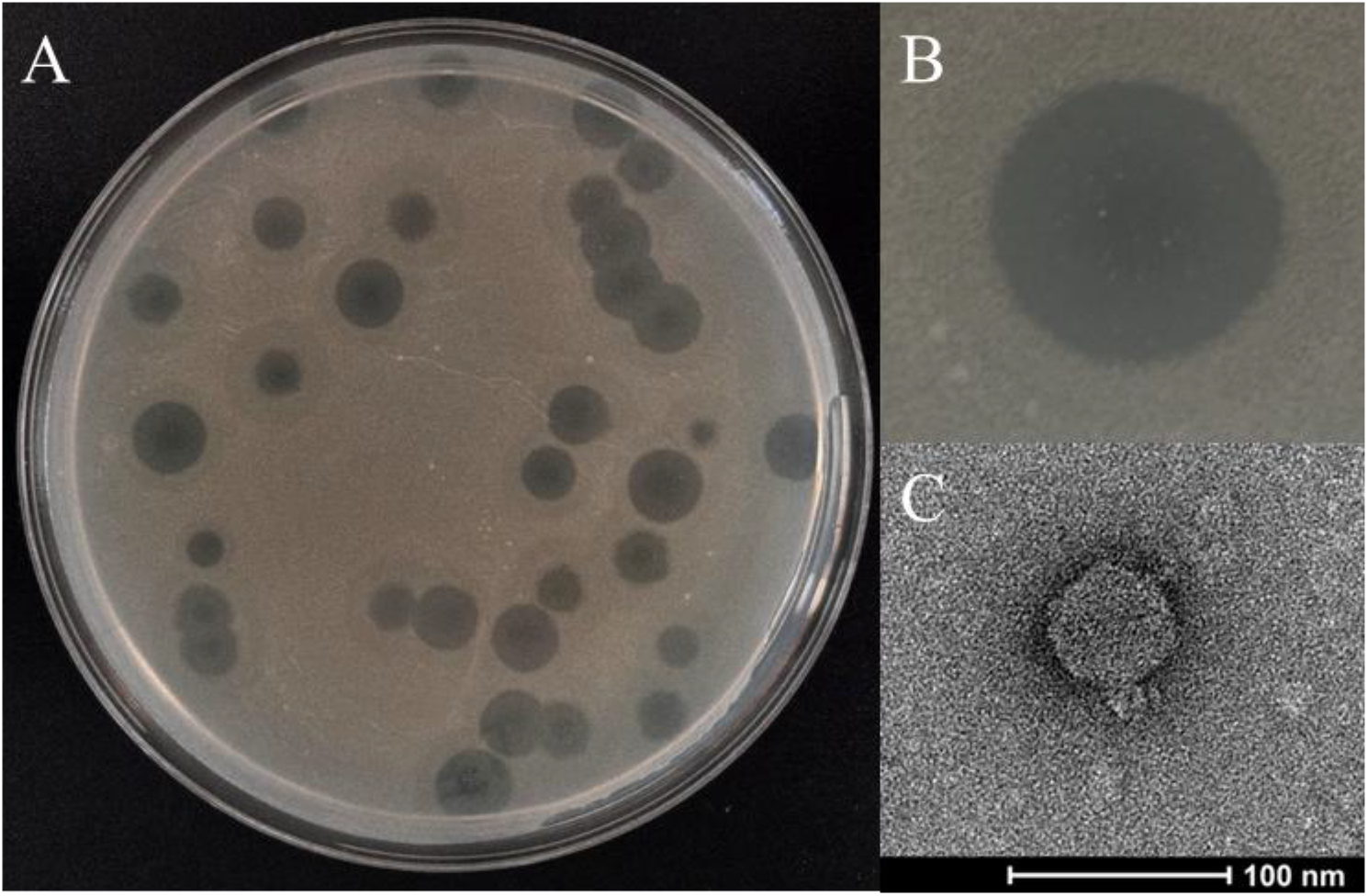
Characterization of phage 17M_Ec17_D. Clear plaques with a semi-transparent halo on overlayer agar (A, B); transmission electron micrograph showing an icosahedral head and short tail, characteristic of the family *Podoviridae* (C). Scale bar, 100 nm.

**Fig. 3.**
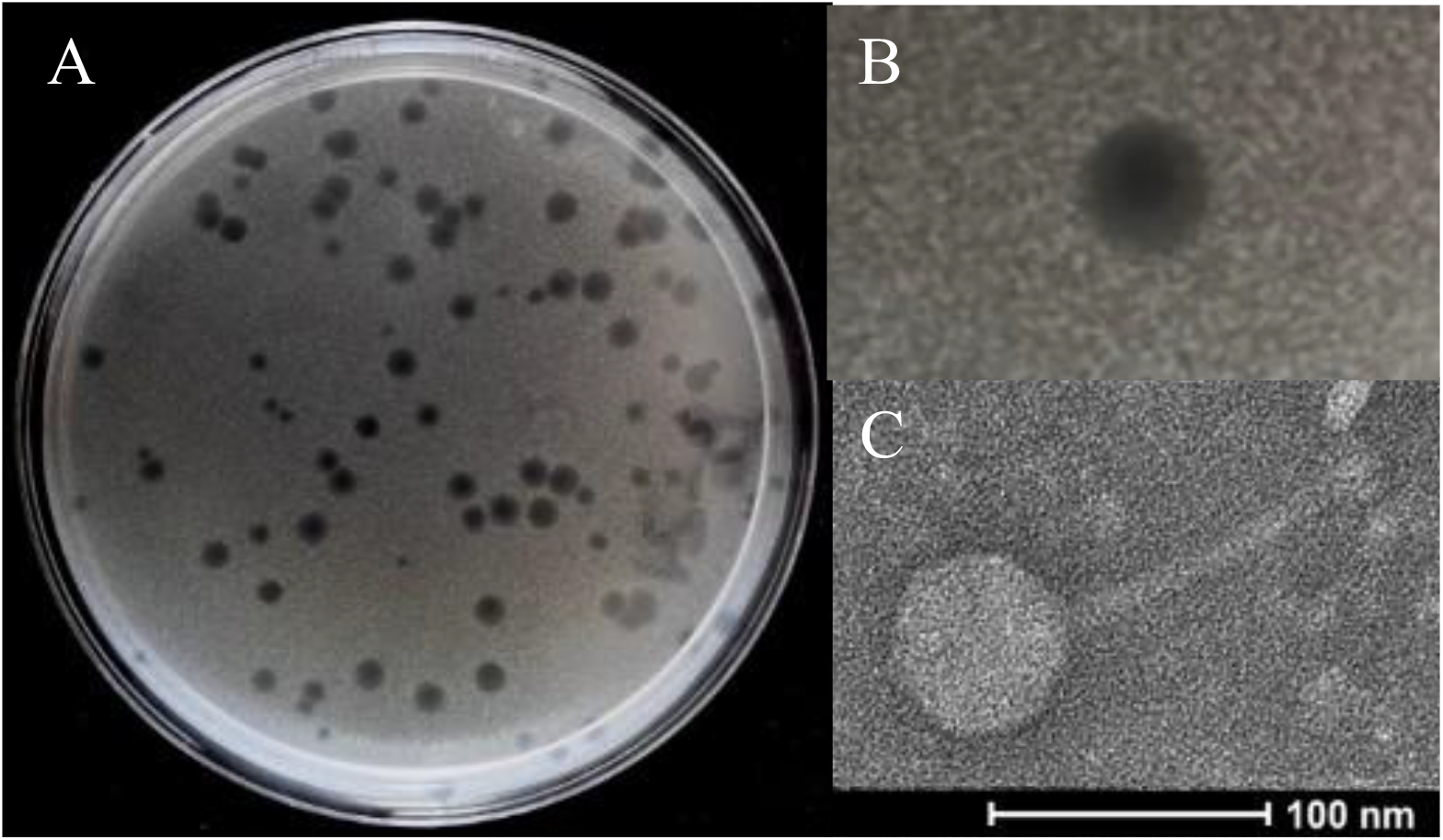
Characterization of phage 22C_Ec22_D. Clear plaques on overlayer agar (A, B); transmission electron micrograph showing an icosahedral head and long non-contractile tail, characteristic of the family *Siphoviridae* (C). Scale bar, 100 nm.

Based on transmission electron microscopy (TEM), 17M_Ec17_D was classified within the family *Podoviridae*, displaying an icosahedral head and a short tail (Fig. 2C), while 22C_Ec22_D displayed *Siphoviridae* morphology, with an icosahedral head and a long, non-contractile tail (Fig. 3C).

### 3.2 WGS Analysis of isolated phages

Genomic analysis enabled species-level classification and according to taxMyPhage and ICTV nomenclature, both phages are candidates for novel species. The closest related species to 17M_Ec17_D was within the genus *Kayfunavirus*, with 89.2% similarity, and for 22C_Ec22_D, it was within the genus *Kagunavirus*, with 71.4% similarity. Nevertheless, further analyses employing at least two independent classification methods will be required for a final and precise taxonomic assignment. The classification, based on head-neck-tail modules using Virfam, confirmed the TEM results, assigning 17M_Ec17_D to the *Podoviridae* Type 3 category and 22C_Ec22_D to the *Siphoviridae* Type 1 (cluster 5) category.

Functional annotation with Pharokka revealed the absence of antibiotic resistance genes, virulence factor genes, or toxin genes in both phage genomes, confirming their safety profile. Genes encoding structural and lysis-associated proteins were identified, including spanins and holins, which are hallmark components of the lytic cycle responsible for host cell lysis and phage particle release. The presence of these genes provides additional molecular evidence of a strictly lytic lifestyle for both phages. Phage lifestyle was further corroborated using BACPHLIP, which searches for lysogenic modules such as integrase, recombinase, and excisionase, none of which were detected in either genome.

The absence of lysogeny modules, combined with the presence of lysis genes, strongly support the classification of both 17M_Ec17_D and 22C_Ec22_D as obligately lytic phages, a desirable characteristic for phage therapy applications. The functional genome organization of both phages is illustrated in Fig. 4.

**Fig. 4.**
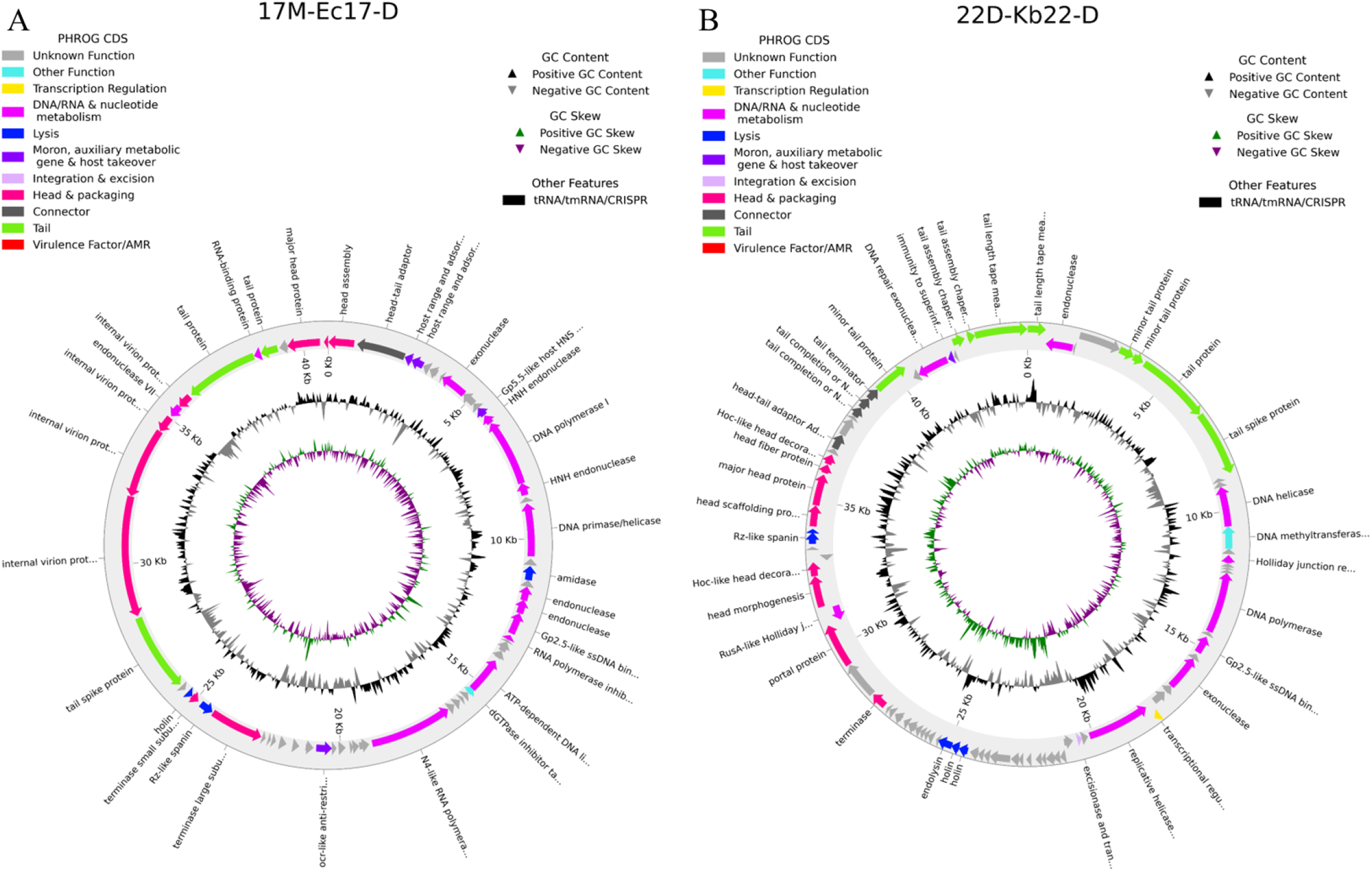
Functional genome annotation of phages 17M_Ec17_D and 22C_Ec22_D. Genome maps with predicted open reading frames color-coded by functional category.

## 4. Discussion

The isolation of phages belonging to the genera *Kagunavirus* and *Kayfunavirus* from urban surface water highlights the critical role of environmental surveillance in combating antimicrobial resistance. These findings align with recent international efforts to source and characterize bacteriophages against difficult-to-treat pathogens, including extensive work targeting drug-resistant *K. pneumoniae* in conflict zones (Green et al., 2025).

The narrow host range of 22C_Ec22_D is consistent with other *Kagunavirus* phages isolated from surface water by Markusková et al., which exhibited lytic activity against only 8% and 3% of tested strains (Markusková et al., 2024). Additionally, Parra et al. reported lytic activity against 9.8% of tested strains for an isolated *Kagunavirus* (Parra et al., 2026). These findings corroborate our results and may indicate that a narrow host range is a characteristic feature of this genus.

Regarding *Kayfunavirus*, the narrow host range observed for 17M_Ec17_D (13%) is consistent with findings reported by Sun et al., who observed similar restrictions against highly virulent *E. coli* and *Salmonella* strains (Sun et al., 2023). According to George et al., *Kayfunavirus* is one of the most abundant phage genera in water environments, yet exhibits limited activity against MDR strains, corroborating our findings (George et al., 2022).

Nevertheless, our host range assays involved a limited number of MDR isolates, and future studies testing a larger panel of strains will be necessary to validate the aspects mentioned above. From a therapeutic perspective, the strict lytic nature of these isolates, confirmed by the presence of holin and spanin genes and the absence of lysogenic elements, validates their safety for potential clinical use (Dicks & Vermeulen, 2024). The observed halos suggest possible depolymerase activity, which could enhance biofilm penetration – a feature worth to further investigate. The clinical value of such tailored treatments is increasingly evident. A recent large-scale study reported clinical improvement in over 77% of complex cases using personalized phage therapy (Pirnay et al., 2024). In this context, the lytic phages characterized here represent valuable additions to therapeutic libraries aimed at combating multidrug-resistant infections.

## 5. Conclusion

In conclusion, urban river water represents a promising reservoir for bacteriophages with strong lytic activity against MDR *E. coli*. The characterized phages, belonging to the genera *Kagunavirus* and *Kayfunavirus*, lack virulence factors, toxin genes, and antibiotic resistance genes, highlighting their potential as safe therapeutic candidates.

To the best of our knowledge, this is the first study characterizing lytic bacteriophages targeting MDR bacteria isolated in Romania, contributing to the growing body of research supporting phage therapy as a viable strategy against antimicrobial resistance.

